# Glacial Legacies – How refugial dynamics shaped the evolution of the Alpine endemic bush-cricket *Anonconotus italoaustriacus*

**DOI:** 10.1101/2025.07.21.665868

**Authors:** Philipp Kirschner, Petra Kranebitter

## Abstract

Mountain ranges like the European Alps harbor large endemic biodiversity shaped by Pleistocene climatic oscillations. The flightless bush-cricket *Anonconotus italoaustriacus*, endemic to the Southern Limestone Alps (SLA) and the eastern Central Alps (CA), provides an ideal model to study the evolutionary and refugial dynamics of endemic alpine arthropods. Using genomic SNP data, we employed phylogenetic analyses, Bayesian clustering, and demographic modeling to investigate the species’ evolutionary history and its refugial dynamics. Our results support a scenario of survival in multiple peripheral refugia during the Last Glacial Period (LGP; 115–11.7 ka), with populations in the southern SLA and eastern CA exhibiting the highest private allelic richness. Postglacial recolonization of interior alpine regions occurred exclusively from refugia on the southeastern margin of the SLA, most likely facilitated by open habitat corridors along transversal valleys. In contrast, populations in the southern SLA exhibited long-term isolation and distributional stasis, emphasizing the importance of small, stable refugia in preserving genetic diversity. We propose that polyandry, a reproductive strategy of *A. italoaustriacus*, contributed to its resilience by maintaining high genetic diversity despite repeated bottlenecks and habitat fragmentation. These findings highlight the importance of integrating evolutionary history into conservation strategies, particularly for alpine endemics with fragmented distributions. Protecting both long-term stable refugia and dynamic evolutionary hotspots is critical for the conservation of *A. italoaustriacus* and other high-altitude arthropods in the face of ongoing environmental change.

## 1 Introduction

Quaternary climate oscillations caused recurring and large scale glaciations of European mountain ranges (Penck & Brückner, 1909; Seguinot et al., 2018). This had a profound impact on the evolution of alpine biota, given that alpine habitats (i.e. habitats above the treeline) were repeatedly covered by ice and became unsuitable during Pleistocene cold-stages such as the Last Glacial Period (LGP; 115–11.7 ka). Alpine biota were forced to shift to suitable refugia whenever ice masses advanced to evade extinction. Permanently unglaciated mountain chains and foothills on the Alpine periphery, so called peripheral refugia (Chodat & Pampanini 1902), were shown to be key for long-term survival of alpine species (Schönswetter et al., 2005). As applicable to mountains in general, the topographical heterogeneity of such peripheral refugia offered suitable niches for species with various ecological preferences and enabled long-term survival (Rota et al., 2024).

During cold-stages, such peripheral refugia were repeatedly delimited from one another by huge glacial streams that ran along large valleys and extended far into the Alpine forelands (e.g. the Etsch/Adige glacier in the Southern Alps; Figure 1). The role of these glacial streams as biogeographic barriers is reflected in the phylogeographies of many Alpine species (Bonato et al., 2018; Hausdorf & Xu, 2023; Nägele & Hausdorf, 2015; Rota et al., 2024; Tribsch & Schönswetter, 2003; Wachter et al., 2016), and also in their extant range limits (Thiel-Egenter et al., 2011). Unsurprisingly, these dynamics of recurring range shifts and recurrent isolation have facilitated large-scale Pleistocene speciation and lineage formation in Alpine biota (Wootton et al., 2025).

**Figure 1.**
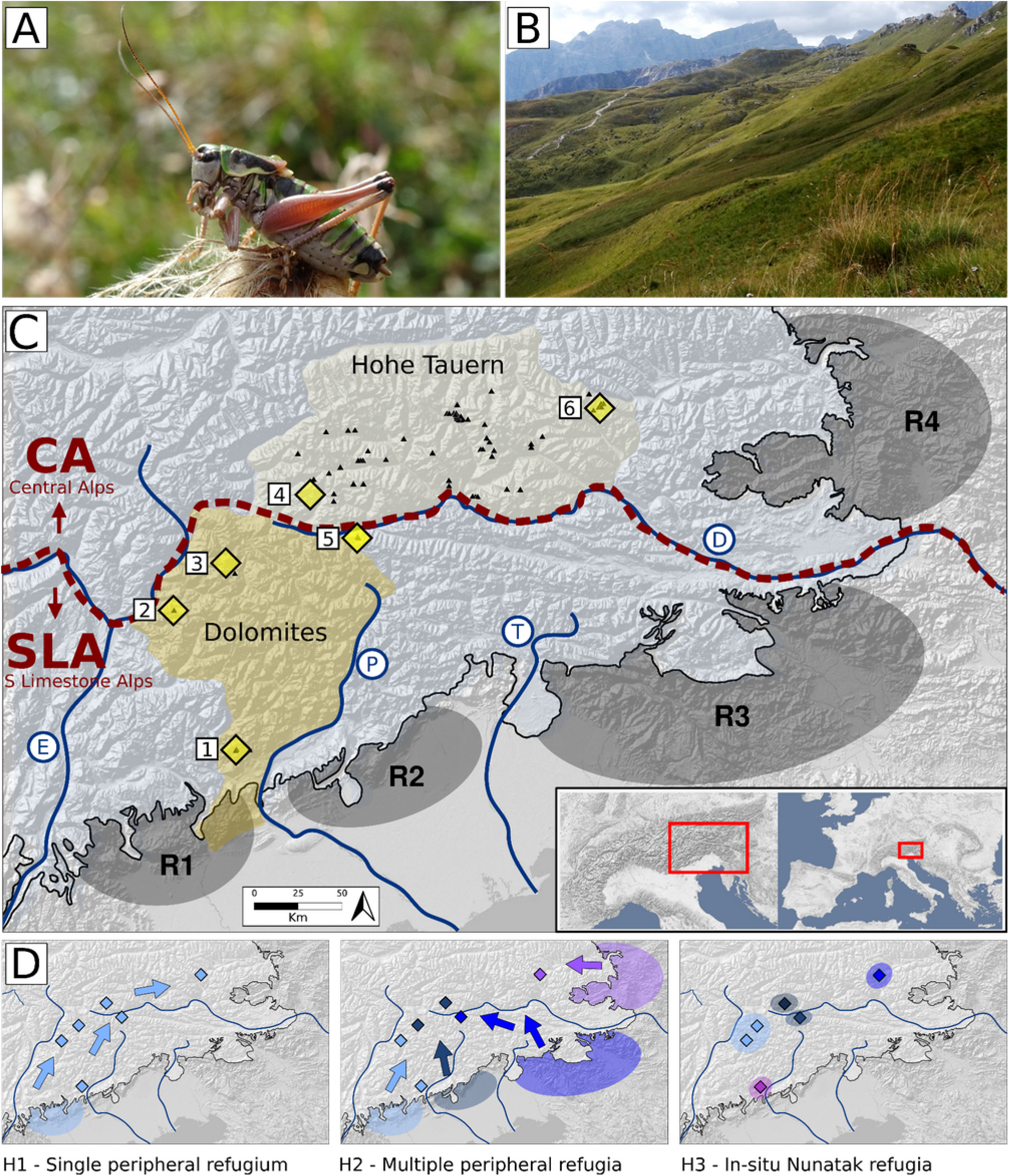
Introduction to the endemic bush-cricket species *Anonconotus italoaustriacus*, including its habitat, current distribution range, and hypotheses regarding its refugial dynamics. **A** An adult male individual of *Anonconotus italoaustriacus* from Vette Feltrine, Southern Dolomites, Italy (Photo credits: Kirschner P. & Kranebitter P.). **B** Typical habitat of *A*. *italoaustriacus*; alpine grasslands southeast of Peitlerscharte/Forcella de Putia, Northern Dolomites, Italy Photo credits: Kirschner D. & Kirschner P.). **C** Map of the Southern Limestone Alps (SLA) and the Eastern Central Alps (CA); geographical entities are delimited by the dashed line and following Grassler (1984). The extant occurrences of *A*. *italoaustriacus* within the Dolomites and the Hohe Tauern mountain ranges (in shades of yellow) are depicted by black triangles; occurrence data retrieved from The Global Biodiversity Information Facility (doi.org/10.15468/dl.nzfnn7) and Galvagni & Fontana, 2004; Zuna-Kratky et al., 2017). Populations sampled for this study are indicated by larger yellow diamonds (1: ‘Vette’, 2: ‘Schlern’, 3: ‘Peitler’, 4: ‘Gsies’, 5: ‘Helm’, 6: ‘Poella’) . The area shaded in blue depicts the maximum glacial extent during the Last Glacial Maximum (LGM; Ivy-Ochs et al., 2009). Relevant major river/glacier systems are highlighted by letters (E: Etsch/Adige, P: Piave, T: Tagliamento, D: Drau/Drava). Grey polygons R1–R4 roughly indicate known refugia for Alpine species (R1: Southern Dolomites, R2: Southern Carnic Alps, R3: Southeasternmost Limestone Alps, R4: Easternmost Central Alps; modified from Tribsch & Schönswetter, 2003). **D** Schematic depiction of three non-exclusive refugial and postglacial expansion hypotheses for *A*. *italoaustriacus*; polygons indicate potential refugial areas, arrows in the same color indicate assumed directionality of postglacial expansion (H1: Single refugium in the Southern Dolomites, H2: Multiple refugia; H3: in-situ survival on Nunataks)

Temperate mountain ranges such as the European Alps are known to harbor a large array of endemic species (Rahbek et al., 2019). Disentangling patterns of endemism has been a longstanding topic in Alpine Biogeography and has contributed significantly to our understanding of evolution and refugial dynamics in Alpine high-altitude biota (Marcuzzi, 1956; Merxmüller, 1952, 1953, 1954). Range-wide studies targeting the vascular plant flora of the Eastern Alps, for example, showed that endemics are not equally distributed within the Eastern Alps, but aggregated in areas associated with refugia (Tribsch, 2004; Tribsch & Schönswetter, 2003). In the Eastern Alps, such aggregations of endemic species are found in the Southern Limestone Alps (SLA, Figure 1), and at the eastern margin of the Alps, specifically the easternmost Central Alps (CA, Figure 1) and the Northern Limestone Alps (Tribsch, 2004). This has been linked to the availability of large and accessible peripheral refugia in close vicinity of the above-mentioned areas (Tribsch, 2004; Tribsch & Schönswetter, 2003). Interestingly, patterns of endemism in ecologically important and species-rich groups, such as Alpine arthropods, are comparably understudied, and knowledge on their spatiotemporal diversification and their refugial dynamics is scarce.

Late Pleistocene refugial dynamics of Alpine mountain arthropods were largely similar to those described for Alpine vascular plants. That is, in short, persistence of populations in peripheral refugia during glacial maxima, followed by subsequent postglacial recolonisation of previously glaciated areas in the interior of the Alps and at higher elevations (Muster, 2000; Wachter et al., 2016). Some authors also suggested long-term refugial survival of arthropods on nunataks (i.e. ice-free mountain tops that protruded the ice shield also during glacial maxima; Holdhaus, 1954; Janetschek, 1956). Such nunataks were previously divided into i) peripheral nunataks in close vicinity to the margin of the ice shield that have not been surrounded by ice at all times, and ii) interior nunataks on the interior of the Alpine ice-shield and permanently surrounded by ice (Holderegger & Thiel-Egenter, 2009). In contrast to the broad evidence supporting survival of vascular plants on these types of nunataks (Rosa et al., 2025; Schneeweiss & Schönswetter, 2011), evidence is scarce and less conclusive for arthropods. Long-term survival on peripheral Nunataks has, for example, been made evident by explicit models based on genetic data for *Trechus* beetles (Lohse et al., 2011) and by ecological niche models for *Megabunus* harvestmen (Wachter et al., 2016). However, survival on non-peripheral Nunataks, a more controversial scenario for arthropods, seems only plausible for specialists of vegetation-free and rocky high-alpine environments such as species from the bristletail genus *Machilis* (Dejaco et al., 2016; Wachter et al., 2012).

The bush cricket *A*. *italoaustriacus* Nadig, 1987 is endemic to the SLA and the eastern CA (Galvagni & Fontana, 2004), and is an excellent model to study diversification and refugial dynamics of an Alpine arthropod in an Alpine hotspot of endemism, and the role of late Pleistocene glaciations on formation of endemism in the area. The species’ range in the SLA is island-like and restricted to a few massifs within the Dolomites and a single, immediately adjacent mountain (Helm/Monte Elmo) in the Carnic Alps (Figure 1; Galvagni & Fontana, 2004). In the eastern CA, the species occurs only in the Hohe Tauern range (Zuna-Kratky et al., 2017), where its distribution range is less island-like with larger and more continuous habitats. *A*. *italoaustriacus* is an alpine species and occurs exclusively above the treeline, where it inhabits open grasslands sometimes with sparse dwarf-shrub cover (e.g. *Juniperus communis*, *Rhododendron* sp.). The species is large and flightless in all life stages, and non-suitable habitats such as forests or shrubland likely act as strong barriers as demonstrated for other Tettigonidae (Kim et al., 2024). Through the lens of this alpine species – its extant range consists mainly of sky islands surrounded by an impassable sea of forest, which renders the species an excellent model to study the impact of recurring glaciations on lineage formation. As all species within the genus *Anonconotus*, *A*. *italoaustriacus* is polyandrous whereas mating is coercive and might involve cryptic female mate choice (Vahed, 2002; Vahed & Carron, 2008).

*A. italoaustriacus* has been listed as endangered by the IUCN because of its strict habitat preference, its narrow range, and the island-like character of many populations (Zuna-Kratky et al., 2017). Timely conservation and management strategies of a species should incorporate evolutionary history of species and populations by disentangling intraspecific genetic variation and by delimiting conservation-relevant entities (Hoban et al., 2022). Ideally, such approaches integrate neutral genetic diversity (Kardos et al., 2021), target the level of genetic connectivity between populations, especially of very isolated occurrences, a population’s demography, and the whereabouts of potential refugia. The latter is particularly important for conservation of any Alpine species, given that refugial populations are genetically most diverse, are the source of present-day genetic variation, and pose evolutionary cradles (Mosblech et al., 2011; Tribsch, 2004).

Here we compare three non-exclusive evolutionary hypotheses for *A*. *italoaustriacus* to explain its extant distribution (Figure 1). We do so by employing Bayesian clustering analyses, phylogenetic analyses, including time-calibrated phylogenies, and demographic modeling, all informed by genomic SNP data. The first hypothesis captures a scenario in which *A*. *italoaustriacus* colonized its present range from a single peripheral cold-stage refugium on the southern margin of the SLA (H1, Figure 1). This scenario implies that the southernmost populations in the Southern Dolomites are phylogenetic sisters to an ingroup containing all remaining populations, and that split ages between ingroup populations should become gradually younger towards the north, reflecting a postglacial expansion from a southwestern refugium to the northeast. The second hypothesis captures a scenario, in which the species has had multiple peripheral cold-stage refugia in the southern and southeastern periphery of the Alps from where the interior of the Alps were colonised postglacially (H2, Figure 1). This scenario implies that some populations in the interior of the Alps are phylogenetic sisters to populations from the eastern margin of the species’ distribution range, and that corresponding splits predate the LGM. In case of a hybrid origin of interior populations, this should be reflected by admixture patterns, and by demographic models. To explain the species present day distribution, both hypotheses imply a fast and large-scale range expansion of the species after the LGM, and colonization of newly emerging habitats before forests expanded into the interior of the Alps. Finally, the third hypothesis captures in-situ refugial survival of the species on nunataks (H3, Figure 1.). Recurring temporal isolation in Nunatak refugia might have played a role and would not stand in contrast to both of the above hypotheses. Such a hypothesis might be backed by deep phylogenetic division between spatially close populations in the interior of the Alps that predate the LGP. Given that refugial areas are reservoirs of genetic diversity, the type and whereabouts of refugia is of large relevance for conservation of alpine species, and results will consequently be interpreted in regard to future conservation and management of the species.

## 2 Methods

### Sampling

A total of 31 specimens from six populations were sampled from the field in the years 2018 to 2024 (Figure 1, Table 1). The sampling covers the species’ entire known distribution range, and includes the southernmost and northeastern margin of its known range. All individuals were collected by hand and transferred to containers containing 96% ethanol. After field sampling, all samples were immediately transferred to the Museum of Nature South Tyrol (Bozen/Bolzano, Italy) for further processing and long-term storage at -20°C.

**Table 1.** Summary of population specific collection data and population summary statistics.

| Population ID | Locality | Collection data |  |  |  | Population summary statistics<br>(based on 1470 variant sites) |  |  |  |  |
| --- | --- | --- | --- | --- | --- | --- | --- | --- | --- | --- |
|  |  | Mountain range | n | NCBI short read archive accession numbers | Coordinates | n private alleles | Pi | Obs. het. | Exp. het. | F<sub>IS</sub> |
| Peitler | Slopes SE of Peitlerscharte | Dolomites (North) | 5 | *** | 46.648N, 11.817E | 127 | 0.159 | 0.210 | 0.134 | -0.086 |
| Schlern | Schlern plateau | Dolomites (North) | 6 | *** | 46.511N, 11.587E | 65 | 0.139 | 0.202 | 0.122 | -0.11 |
| Helm | S slopes of Helm | Carnic Alps | 5 | *** | 46.712N, 12.384E | 105 | 0.149 | 0.217 | 0.132 | -0.120 |
| Gsies | Slopes E of Kapairalm (Gsies) | Hohe Tauern | 5 | *** | 46.845N, 12.188E | 117 | 0.161 | 0.224 | 0.142 | -0.112 |
| Poella | Slopes E of Lanisch-Ochsenhuetten (Poellatal) | Hohe Tauern | 5 | *** | 47.069N, 13.454E | 160 | 0.164 | 0.229 | 0.145 | -0.115 |
| Vette | Plains of Busadelle Vette | Dolomites (South) | 6 | *** | 46.094N, 11.842E | 410 | 0.203 | 0.254 | 0.179 | -0.082 |

### Sequencing

DNA extraction, sequencing library preparation and DNA sequencing was done by Biosearch Technologies (LGC Genomics, Berlin, Germany). Genomic DNA was extracted from femoral tissue and sequenced using genotype by sequencing (GBS; modified from Elshire et al. 2011). For the employed GBS the restriction enzyme MSII was used to fragment genomic DNA, followed by a normalisation step in which abundant fragments were reduced prior to sequencing. Sequencing was done using the Illumina sequencing protocol and an Illumina NovaSeq platform (Illumina, San Diego, USA) aiming for 150 bp paired-end reads.

### Raw sequence processing and SNP calling

Raw read data was checked for presence of cutsites (--e mseI), quality filtered (--q), and truncated to a length of 140 bp (-t 140) using *process_radtags* included in Stacks 2.68 (Catchen et al., 2013). Loci were assembled from primary and secondary reads via denovo_map using the parameters -m 5 (minimum coverage), -M 3 (maximum mismatches between loci within individuals) and -n 4 (maximum mismatches between loci of different individuals; Catchen et al., 2013). These parameters were inferred from a subset of the data containing a randomly selected individual per population following an initial parameter optimization (Paris et al., 2017). Population summary statistics (n of private alleles, nucleotide diversity, observed and expected heterozygosity, inbreeding coefficient F_IS_) were calculated for each population using *populations* using SNPs that were present in at least 80 percent of individuals (-R 0.8), and using a single SNP per fragment (--write-single-snp; Catchen et al., 2013). F_ST_ values were calculated using the same settings. Isolation by distance (IBD) was tested based on transformed F_ST_ (F_ST_/1-F_ST_) and log-transformed euclidean distances using the gl.ibd function implemented in dartR (Gruber et al., 2018; Mijangos et al., 2022).

### Bayesian Clustering

For Bayesian clustering in STRUCTURE (Pritchard et al., 2000) analysis SNPs were exported from the loci catalog to STRUCTURE format (--structure) including only SNPs that were at least present in 65% of all individuals (-R 0.65), whereas only a single SNP per fragment was used to avoid including linked SNPs (Catchen et al., 2013); the final STRUCTURE input files contained 31 individuals and 9463 SNPs. Clustering analysis was done using the admixture model in STRUCTURE for K=1 to 5 clusters running 10 replicates per K for 1,000,000 generations and discarding 200,000 generations as burnin. STRUCTURE results were summarized using CLUMPAK (Kopelman et al., 2015). The most likely number of K was evaluated using the Delta K criterion (Evanno et al., 2005).

### Phylogenomic analyses and divergence time estimation

A phylogenetic network was created using the software Splitstree 4 (Huson & Bryant, 2006) using uncorrected p-distances. Input data contained only SNPs present in at least 75% of all individuals, and SNPs with a minimum mean coverage of 5 and was prepared using populations (Catchen et al., 2013) and vcftools (Danecek et al., 2011). The resulting vcf file included 5717 SNPs and was transformed to nexus format using the software vcf2phylip v. 2.6 (Ortiz, 2019).

Phylogenetic tree reconstruction and divergence time estimations were done using the SNAPP module (Bryant et al., 2012) integrated in Beast 2.7.8 (Bouckaert et al., 2014). To reduce computational load, only two or three individuals per population were included for the analyses, whereas individuals with less missing data were preferred. *A*. *alpinus* was used as an outgroup and sequences of three individuals of *A*. *alpinus* from two localities (1 specimen: Col du Lautaret, France, 45.0383N, 6.3956E; 2 specimens: Morcles, Switzerland, 46.2153N, 7.0618E), including the species’ type locality (Yersin, 1858), were selected. Input data was prepared using the script snapp_prep.rb (available at: https://github.com/mmatschiner/snapp_prep) using a vcf file that included only a single SNP per fragment and only biallelic SNP (in total 3840 SNPs). The median stem age of *A. alpinus* and *A. italoaustriacus* was used as a calibration point and set to 1.87 Ma using a lognormal prior (mean = 1.95, standard deviation = 0.28). This estimate was derived from a rate calibrated phylogeny based on sequences from two mitochondrial genes (16S, *cytochrome b*) and including all extant *Anonconotus* species (Kirschner et al., in prep.). Four independent SNAPP runs were started and each MCMC chain was run for at least 3,000,000 generations. Log files from each SNAPP run were inspected in Tracer 1.7 (Rambaut et al., 2018) to ensure sufficient effective sample size (> 200), stationarity of the MCMC chain for all parameter estimates, and convergence between chains. Trees were summarized into a maximum clade credibility tree using mean common ancestor node ages in treeannotator 2.7.3 (Bouckaert et al., 2014). To independently verify node age estimates, an additional phylogenetic dating analysis was done using four mitochondrial genes from 24 individuals of *A*. *italoaustriacus* and including *A*. *alpinus* as an outgroup (Martinez et al. 2022; details in Supplementary Material). This was also done as the analysis in the original paper lacked some information, e.g. on the range of age estimates.

### Demographic analyses

Demographic models were fitted on the allele frequency spectrum using the software dadi (Gutenkunst et al., 2009). This was done using the *dadi pipeline* (Portik et al., 2017), that allows to fit a large set of predefined models representing common evolutionary scenarios. Here, a two-step hierarchical approach was employed. In the first step, a set of three population models that were designed to capture migration and population size changes at different time horizons were used. Specifically, these models assume one population (Pop 1) to be the sister to two diverging populations (Pop 2 and Pop 3), whereas Pop 3 is considered to be located geographically in between (Firneno et al., 2020; Figure S1). This set of models allows testing between different refugial scenarios, i.e. if gene flow between isolated populations occurred ancestrally, prior to divergence, or via secondary contact, after population divergence, and if this gene flow occurred unidirectionally or symmetrically (Firneno et al 2020). Two additional models that captured secondary gene flow between Pop 2 and Pop 3 were added (unidirectional and symmetrical; Figure S1). Pop 1–3 were defined based on a priori information from phylogenetic analyses and Bayesian clustering (see results section; Figure 2), i.e. as ‘Southern Group’ (Pop 1), ‘Eastern Group’ (Pop 2) and ‘Interior Group’ (Pop 3); the latter was defined as the geographically in-between population for modelling. In a second step, so-called ‘island models’ were employed. These models capture divergence scenarios between two populations that include either founder events or vicariance (Figure S1; Charles et al., 2018). Here, these models were used for the ‘Eastern Group’ and the ‘Interior Group’, whereas the latter was defined to be derived from the first based on the a priori information from phylogenetic analyses (Figure 2, 3).

**Figure 2.**
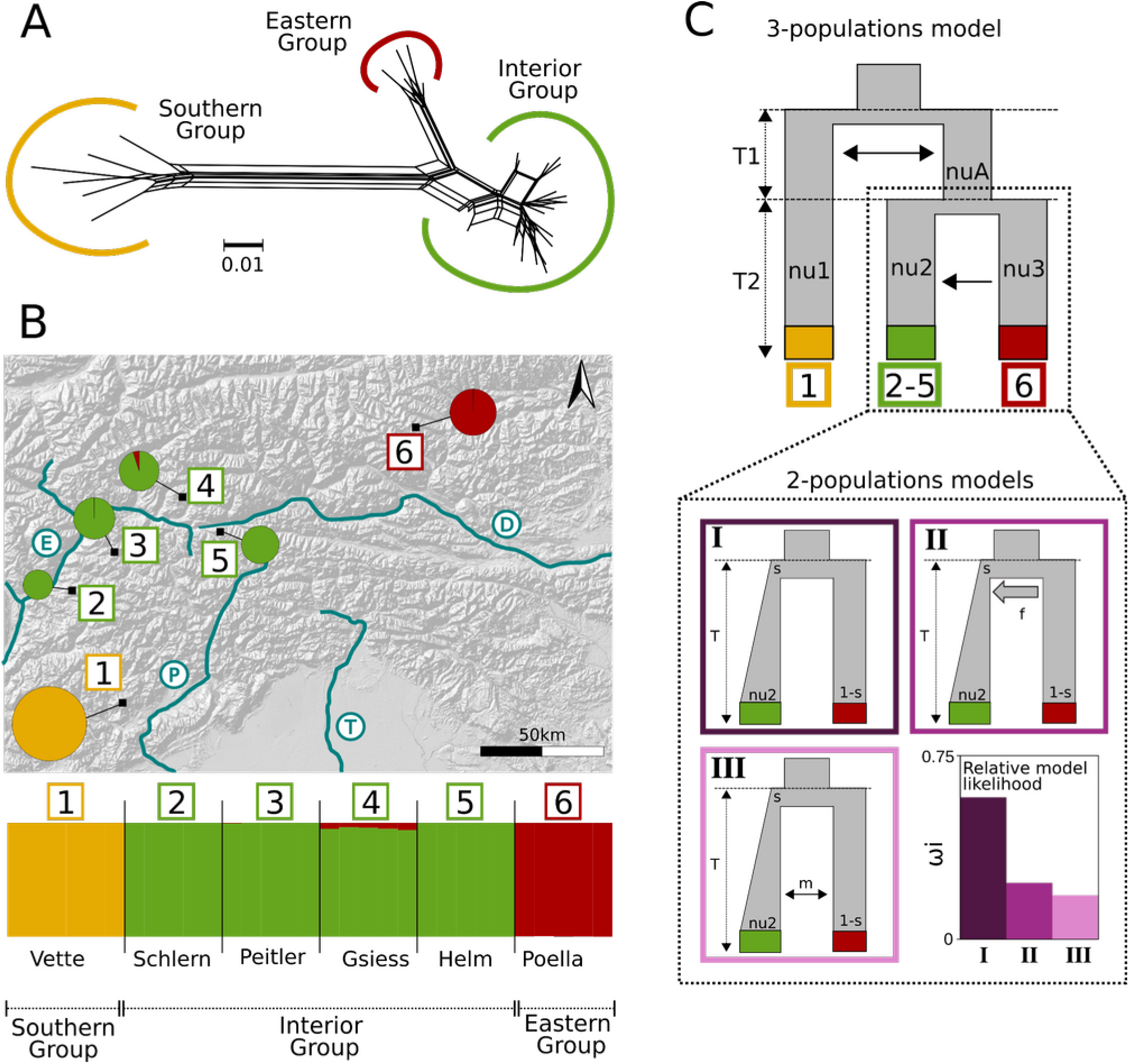
Phylogenetic relationships, distribution of genetic variation and demographic history of *A*. *italoaustriacus* based on genomic SNP data. Group definition (‘Southern Group’, ‘Eastern Group’, ‘Interior Group’) and population color-coding are the same in subfigures A–C. **A** Phylogenetic network based on p-distances. **B** Proportion of shared genetic variation inferred via Bayesian clustering for each population (pie charts in map) and for each individual (each bar in the plot represents one individual) assuming K=3. Diameters of pie charts correspond to the n of private alleles in each population (n=65– 410; Table 1). **C** Schematic summary of the best performing demographic models. The best performing 3-population model capturing relationships between ‘Southern Group’, ‘Interior Group’ and ‘Eastern Group’ is shown on top (ancestral migration between ‘Southern Group’ and the ancestor of ‘Interior Group’ and ‘Eastern Group’, in the first epoch, followed by unidirectional migration from ‘Eastern Group’ to ‘Interior Group’ in the second epoch). The three best performing 2-population models capturing relationships between ‘Interior Group’ and ‘Eastern Group’ (I: founder event without migration; II: founder event with discrete post-split admixture; III: founder event with subsequent continuous symmetric migration), and a barplot indicating the relative conditional likelihood of each model are shown below. All included models are summarized in Figure S1.

**Figure 3.**
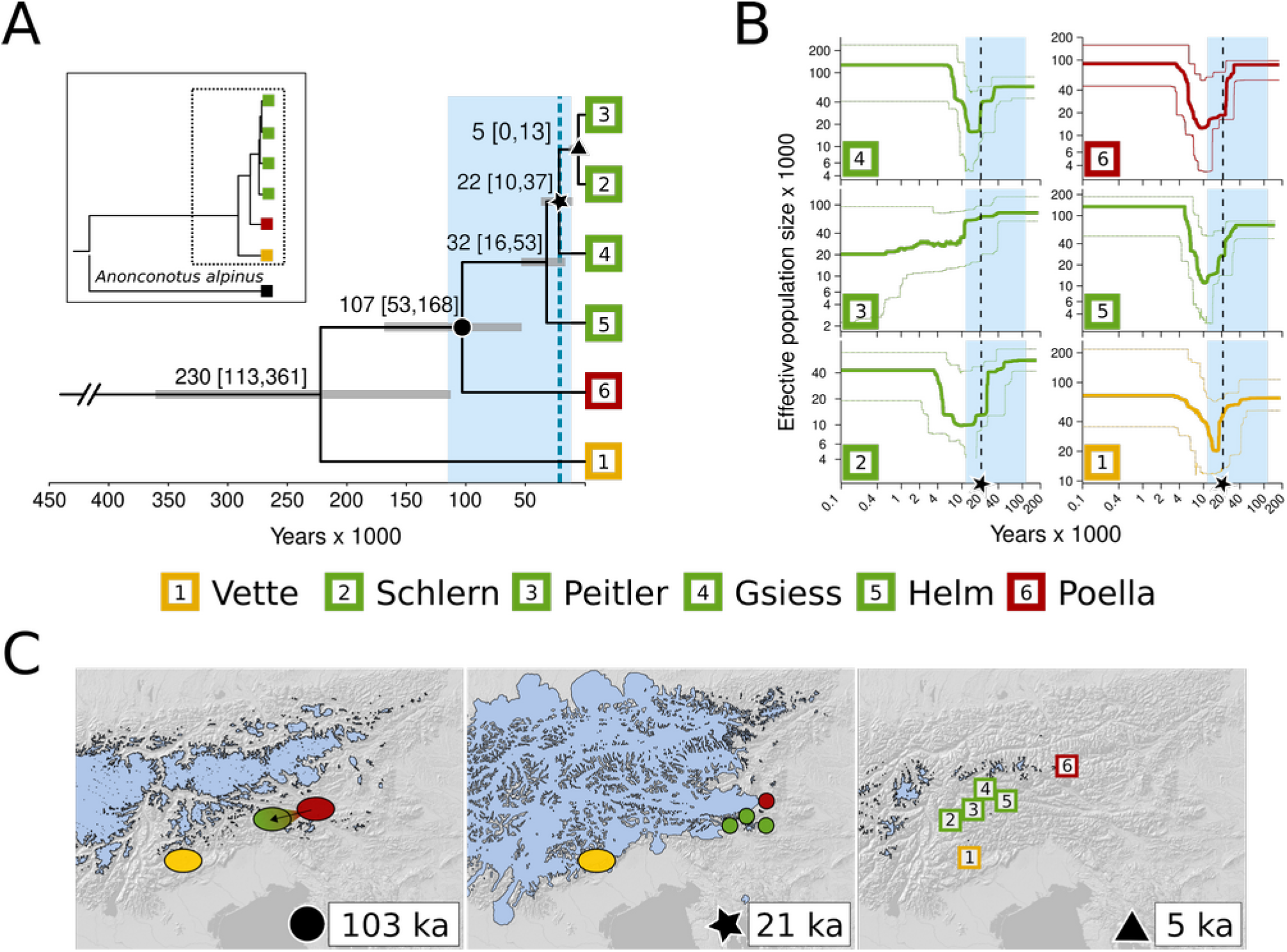
Time-calibrated phylogenetic tree, population-wise demographic trajectories and hypothetical diversification scenarios. **A** Time-calibrated coalescent tree. Node labels indicate the inferred median ages followed. Intervals of 95% highest posterior density are given in parentheses and are represented by grey bars.The area shaded in blue indicates the Last Glacial Period (115–11.7 ka), and the dashed line indicates the Last Glacial Maximum (∼21 ka). The caption shows the split of *A*. *italoaustriacus* and the outgroup *A*. *alpinus* that was used to calibrate the phylogeny. Node symbols correspond with symbology indicating time in subfigures B–C. **B** Stairway plots indicating the fluctuation of effective population size through time for each population. Thick lines indicate mean values, thin lines the 95% confidence intervals; scales are not linear. The area shaded in blue indicates the Last Glacial Period (LGP; 115–11.7 ka), and the dashed line indicates the Last Glacial Maximum (LGM; ∼21 ka). **C** Hypothetical diversification scenario for *A*. *italoaustriacus* at three points time and the corresponding glacial extents (Ivy-Ochs et al., 2009; Seguinot et al., 2018). Polygons roughly indicate the putative location of populations at the given time. The first panel shows distributional stasis of ‘Vette’ and a founder event with ‘Poella’ as source population at the onset of the LGP. The central panel shows refugial isolation of all populations and diversification of ‘Helm’, ‘Gsies’ and the ancestor of ‘Peitler’ and ‘Schlern’ in microrefugia during the LGM. The right panel shows the distribution of extant populations 5 ka which is similar to today’s situation, whereas rectangles correspond to each populations actual present-day location. Color coding and symbols at the bottom correspond with subfigure A.

All employed demographic analyses were calculated on the basis of folded minor allele frequency spectra (AFS). These spectra were generated using the script easySFS.py (github.com/isaacovercast/easySFS) that utilizes functions of the dadi package (Gutenkunst et al., 2009). The underlying vcf files were pre-filtered using vcftools (Danecek et al., 2011), and SNPs missing in more than 70% of individuals (--max-missing) and those with a coverage below 5 (--min-meanDP) were excluded. When necessary, AFS were down projected to increase the number of segregating sites using easySFS.py, resulting in 8, 8 and 8 (‘Southern Group’,‘Interior Group’, ‘Eastern Group’) alleles for the three population models, and 24 and 8 (‘Interior Group’,’Eastern Group’) alleles for the two population island models. The dadi pipeline was run in four rounds using 220 replications per model. The best performing models were selected based on the Akaike information criterion (AIC) and conditional model probabilities were evaluated using Akaike weights (ωi; Burnham & Anderson, 2002).

Past changes of effective population size were modeled for each population using Stairwayplot2 (Liu & Fu, 2020), a software that also utilizes the AFS. AFS were generated in <u>easySFS.py</u> for each population separately and without using downprojection. Underlying vcf files were again pre-filtered in vcftools (Danecek et al., 2011), and SNPs missing in more than 90% of individuals (--max-missing) and those with a coverage below 5 (--min-meanDP) were excluded. To transform the resulting demographic estimates from generations before present into actual geologic time estimates, a mutation rate of 2.8*10^−9^ (Keightley et al., 2014) and a generation time of one year were used. All remaining model parameters were adjusted following the recommendations of the Stairwayplot2 manual (Liu & Fu, 2020).

## 3 Results

### Sequencing and population statistics

The SNP calling in the denovo_map pipeline yielded a total of 1,149,201 loci with a mean length of 231 base pairs and a 9x mean coverage. These loci were composed of 268,251,074 invariant sites and 974,830 variant sites. Population-wise calculations of nucleotide diversity and the amount of private alleles are given in Table 1, proportion and distribution of private alleles in Figure 2. Observed heterozygosity (ObsHet) was larger than the expected heterozygosity (ExpHet) in all populations, suggesting a larger heterozygosity than expected under Hardy Weinberg Equilibrium (Table 1). Inbreeding coefficient F_IS_ was smaller than zero in all (-0.08 to -0.12) in all populations suggesting a lack of inbreeding (Table 1). F_ST_ ranged from 0.11 to 0.35, indicating strong differentiation between extant populations (Figure S2). Testing for IBD resulted in a non-significant positive correlation between F_ST_ and genetic distance (alpha = 0.05, p = 0.13, R² = 0.288).

### Bayesian clustering and admixture analysis

Bayesian clustering analyses identified a maximum of three groups (‘Southern Group’, ‘Interior Group’, ‘Eastern Group’, Figure 2B; consistent in K=3–5, Figure S3) whereas a separation into K=3 clusters was suggested to be optimal (Figure 2). The population ‘Vette’ was resolved as a separate group (‘Southern Group’) in all (K=2-5) analyses, whereas when K=2 was assumed, shared variation with the population ‘Poella’ was found. The population ‘Poella’ was resolved as a separate group (‘Eastern Group’; K=3–5) with a small proportion of shared variation with the ‘Interior Group’ in K=4–5 (Figure S3).

### Phylogenomic analyses and divergence time estimation

Phylogenetic network analyses resolved three groups corresponding to the groups resolved by Bayesian clustering (Figure 2A–B). This was similarly resolved by the coalescent tree inferred via SNAPP. This tree showed shallow divergence between populations from interior parts of the SLA (‘Helm’, ‘Peitler’, ‘Schlern’) and the eastern CA (‘Gsies’, Figure 3A). ‘Poella’ from the CA was resolved as a divergent phylogenetic sister to the aforementioned (Figure 3A). ‘Vette’ from the southern margin of the SLA was resolved as a phylogenetic sister to all other populations (Figure 3A). All resolved nodes were highly credible with posterior probabilities of 1 (not shown). Molecular dating resulted in a mean node age of 1.77 Ma (95% highest posterior density [HPD]: 0.9, 2.76 Ma) for the split between *A*. *italoaustriacus* and the outgroup *A*. *alpinus*. Diversification within *A*. *italoaustriacus* occurred significantly later, and ‘Vette’ split only 230 ka (95% HPD: 114, 361 ka), coinciding the penultimate glaciation and predating the last interglacial (LIG: 115– 130 ka). Separation of ‘Poella’ and ‘Peitler’, ‘Schlern’, ‘Helm’ and ‘Gsies’ occurred 107 ka (95% HPD: 53, 168 ka), or shortly after the beginning of the LGP (115 ka–11.7 ka). Further diversification (32 and 22 ka) coincided roughly with the LGM, and only the populations ‘Peitler’ and ‘Schlern’ diverged postglacially (5 ka). The rate calibrated phylogeny based on four mitochondrial genes resulted in similar divergence time estimates for the split between *A*. *alpinus* and *A*. *italoaustriacus* (1.88 Ma; 95% HPD: 0.95, 2.86 Ma), and the crown diversification age of the ingroup (330 ka; 95% HPD: 167, 531 ka; Figure S6).

### Demographic modeling

Models with ancestral gene flow and unidirectional secondary contact resulted as the best fitting models from the three population modeling (Figure 2C; Table S1). The best performing model captured ancestral migration between Pop 1 ‘Southern Group’ and the ancestor of Pop 3 ‘Interior Group’ and Pop 2 ‘Eastern Group’, and unidirectional migration from Pop 2 to Pop 3 (Figure 2C; Table S1; Figure S4). Founder event models were the best fitting models in the two population modelling (Figure 2C; Table S1; Figure S5). Of the three best performing models, none showed significantly better fit (ΔAIC < 2; Table S1). Conditional probabilities (Figure 2C; Table S1) were largest for the simplest model capturing a founder event without migration (ωi = 0.61), followed by a founder event model with discrete post-split admixture (ωi = 0.22), and a founder event model with subsequent continuous symmetric migration (ωi = 0.15). The s parameter, the fraction of the source population going into the founded population, was ≤ 10% in all three models.

Demographic trajectories of all populations resolved a decline of effective population size coinciding with the LGM (Figure 3B). Effective population sizes recovered after the LGM and reached pre-LGM levels in all populations except population ‘Peitler’.

## 4 Discussion

### LGM survival in multiple refugia in the periphery of the Southern Alps

This study provides detailed insights into the evolution of *Anonconotus italoaustriacus*, an endangered mountain arthropod species endemic to alpine grasslands of the eastern Central Alps (CA) and the Southern Limestone Alps (SLA). We outline a scenario in which *A*. *italoaustriacus* has survived the Last Glacial Period (LGP; 115–11.7 ka) in two distinct peripheral refugial areas on the southern and southeastern margins of the SLA, overlapping areas emphasized as refugia for alpine plant species (R1 and R3, Figure 1, Schönswetter et al., 2005; Tribsch & Schönswetter, 2003). This is reflected by the distribution of private allelic richness, the geographic distribution of genetic variation, and by phylogenetic analyses (Figure 2A, Figure 3A, Table 1). Populations that today occur at the southern margin of the SLA (‘Vette’) and in the the eastern CA (‘Poella’; reflecting both the southern and eastern range limits of the species) exhibited high levels of private allelic richness, were separated into distinct genetic clusters, and appeared as respective phylogenetic sisters to the remaining populations located in interior parts of the Alps (all populations in the ‘Interior Group’; Figure 2A–B, Figure 3A).This evidence supports the a priori outlined ‘multiple refugia hypothesis’ (H2, Figure 1D), and refutes the ‘single southern refugium hypothesis’ (H1, Figure 1D).

In addition no evidence in support of the ‘in-situ survival hypothesis’ (H3, Figure 1D) was found, and we consider such a scenario to be unsubstantiated and misleading. Populations of the ‘Interior Group’ previously considered to be candidates for survival on nunataks exhibited low levels of private allelic richness (Figure 2B, Table 1), contrasting long-term stability, and originated only at the end or after the LGP (Figure 3A). The study that initially raised this “in-situ survival hypothesis” (Martinez-Sañudo et al., 2022), based this scenario on phylogenetic analyses that suggested split ages of 800 ka to 2200 ka between populations that correspond to the populations ‘Vette’, ‘Schlern’, ‘Peitler’, ‘Helm’ and ‘Gsies’ in our study (Figure S6). However, despite discussions with the main authors of Martinez-Sañudo et al. (2022) and a critical reanalysis of their data (Supplementary Material; Figure S6), we were unable to reproduce the reported age estimates. While we made efforts to address this issue collaboratively, the authors were unable to provide a definitive resolution of the issue. We suggest that the original estimates in Martinez-Sañudo et al. (2022) may reflect methodological artifacts, raising concerns about the reliability of their conclusions. Conversely, node age estimates derived from our reanalysis of their data were consistent with those based on our own independently generated data (Kirschner et al., in prep.; Figure 3A; Figure S6).

### Long-term isolation and distributional stasis in the Southern Dolomites and postglacial recolonization of the Alps along transversal valleys

Despite the existence of multiple refugia, postglacial recolonisation of interior parts of the Alps occurred exclusively from the latter (R3, Figure 1C), but not from refugia on the southern margin (R1, R2, Figure 1C). This is evident as all populations in the interior of the Alps form a monophyletic group (‘Interior Group’) that showed no signs of admixture or secondary contact with the ‘Southern Group’ (Figure 2B). Within the ‘Interior Group’, the age of populations decreased gradually with geographic distance from east to west, suggesting a stepping-stone like mode of colonisation. The Initial separation of populations ‘Vette’ (‘Southern Group’) and populations of the ‘Interior Group’ and ‘Eastern Group’ occurred 230 ka (Figure 3A), possibly by fragmentation of a continuous or loosely interconnected range along the southern margin of the SLA (Figure 3C). The best fitting demographic model supports such a scenario as it captures ongoing ancestral gene flow after initial lineage separation (Figure 2B). Separation 230 ka (Figure 3A) coincides with the warm-stage before the penultimate glacial period (∼200–135 ka), and it is possible that advancing ice masses may have triggered this initial separation of lineages. This would fall in line with the diversification dynamics observed in other narrow endemics from the same region, where lineage formation was linked to recurring expansion of large glaciers along the Piave valley (Rota et al., 2024).

The ‘Southern Group’ was found to be confined to the Vette Feltrine range in the Southern Dolomites where it existed in isolation since at least the penultimate glacial period (Figure 2B–C, Figure 3A,C). Similar to other authors (Galvagni & Fontana, 2004; Buzzetti pers. comm., 2024), we found that the species is absent from nearby ranges. We suggest that this hints at long-term distributional stasis (Kropf et al., 2002), likely reinforced by the island-like nature of the Vette Feltrine. Like other ranges along the southern margin of the SLA, the Vette Feltrine are surrounded by deep valleys that have acted as strong dispersal barriers, causing significant phylogenetic divergence between spatially close lineages, especially when dispersal capacity is limited (Hausdorf & Xu, 2023; Nägele & Hausdorf, 2015; Rota et al., 2024). This also applies to *A*. *italoaustriacus*, which is a wingless species that requires continuous open habitats to disperse. We suggest that significant range expansions in this area was limited to comparably short periods, as these valleys were either covered by glaciers (cold periods) or impassable forests (warm periods) during most of the late Quaternary (Figure 4).

**Figure 4.**
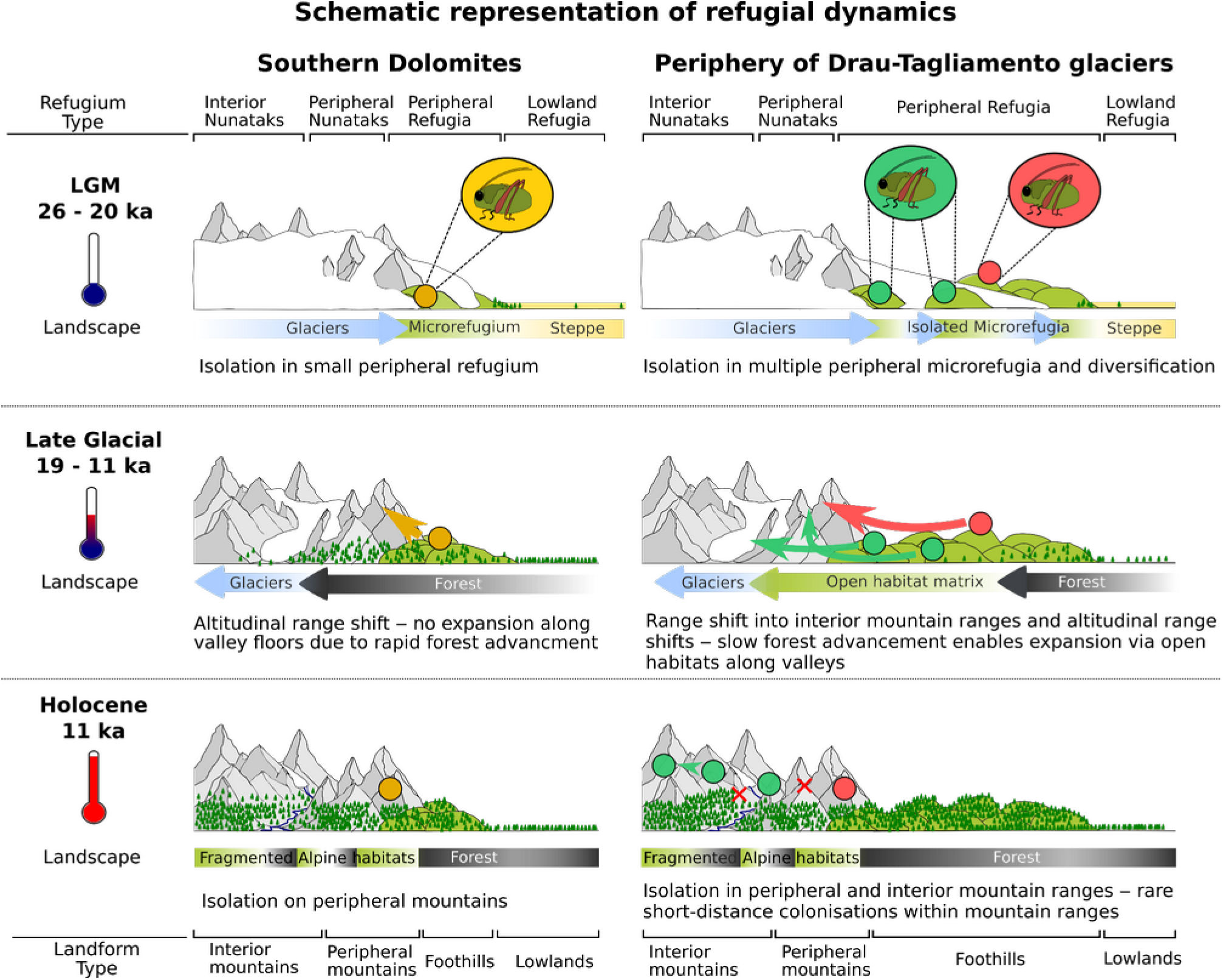
Summary and schematic representation of contrasting refugial scenarios for *A. italoaustriacus* in the Southern Dolomites and the periphery of the Drau-Tagliamento glaciers. Refugia in the Southern Dolomites were smaller and more isolated, with rapid forest expansion occurring immediately after deglaciation. This facilitated altitudinal range shifts for populations of *A. italoaustriacus* from peripheral refugia but hindered their postglacial range expansion into the interior parts of the Alps, as longitudinal valleys were quickly covered by impassable forests. In contrast, microrefugia in the periphery of the Drau-Tagliamento glaciers were larger and more abundant, supported by a local topography characterized by extensive and diverse foothills. Forest expansion following deglaciation was slower due to the influence of a more continental climate. These conditions allowed populations of *A. italoaustriacus* to expand their ranges more freely, facilitating both altitudinal and longitudinal movements into interior alpine regions along open latitudinal valleys. Landscapes dominated by glaciers, forests and steppe are considered to be impassable for *A*. *italoaustriacus*. Grasshopper icons and circles in different colors represent distinct genetic lineages.

We hypothesize that the Vette Feltrine may represent especially suitable refugia for *A*. *italoaustriacus* for two key reasons: first, the range features a large elevational gradient from its foothills to its peaks (400–2500 m) that is not interrupted by significant topographic barriers. This may have enabled elevational range buffering throughout climatic changes even for a species with limited dispersal ability. Second, the range includes extensive alpine plains, for example the ‘Busa delle Vette’, featuring extensive, continuous grasslands (Pignatti & Pignatti, 1983). Such grasslands are critical for the species’ long-term survival, and are not present to such an extent in any neighboring range. We are unable to finally clarify whether certain refugia are particularly well-suited for *A. italoaustriacus* or if the observed patterns are simply the result of stochastic processes. However, we emphasize that this may represent an intriguing avenue for future research. Comparative studies that integrate fine-scale evolutionary data from specialist species could provide fresh insights into these dynamics.

The diversification and range dynamics of populations in the ‘Eastern Group’ and ‘Interior Group’ differed starkly from those of the ‘Southern Group’ and appear to have been considerably more dynamic (Figure 4). The species’ LGM refugia were most likely located in the periphery of the Tagliamento-Drau glacier system, a known refugium for alpine species (R3, Figure 1C; Schönswetter et al., 2005; Tribsch & Schönswetter, 2003). Cold-stage refugia significantly north of the Drau glacier seem implausible regarding the general absence of the species from most of the eastern CA (R4, Figure 1C). Initial diversification between the ‘Eastern Group’ and ‘Interior Group’ occurred at the beginning of the LGP (107 ka, Figure 3A), coinciding with a drop in global temperatures that marked the first large-scale glacial advance of the LGP (Figure 3C; Seguinot et al., 2018). This cooling similarly caused a decline in closed forests, while open habitats with dwarf shrubs— resembling the present-day treeline ecotone—expanded at low and mid-altitudes (Drescher-Schneider, 2000). These newly available suitable habitats may have simultaneously facilitated the range expansion of diverging *A. italoaustriacus* lineages.

Such a scenario is backed by the best-performing demographic models, which uniformly captured a founder event with exponential growth (Figure 2C, Table S1). Divergence between the ‘Eastern Group’ and the ‘Interior Group’ remained upright throughout the LGP without evidence for secondary contact (Figure 2B–C), and diversification continued during the LGM (20–30 ka, Figure 3A, C). We interpret this as evidence for the existence of recurrently isolated microrefugia (Rull, 2010) in the periphery of the Tagliamento-Drau glacier system. Advancing glaciers likely acted as barriers and potentially facilitated allopatric formation of lineages on small spatial scales (Figure 3C, Figure 4; Mosblech et al., 2011). Notably, the Tagliamento-Drau glacier system was particularly dynamic and is known for a two-fold expansion and retreat cycle during the LGM (20–30 ka; Monegato et al., 2007). Eventually this process has additionally favored such a diversification in microrefugia in this area.

We outline a scenario in which the interior parts of the Alps—comprising most of the species’ extant range—were colonized exclusively from southeastern refugia located in the periphery of the Tagliamento-Drau glacier system (R3, Figure 1C). We suggest that this expansion occurred along large transversal valleys, such as the Drautal (indicated as ‘D’, Figures 1C, 2B). This may be already evident by the species’ extant occurrences on mountain ranges flanking these valleys and their tributaries (Figures 1C, 2B), and finds strong support by decreasing split ages and decreasing allelic richness from east to west (Figures 2B, 3A). While transversal valleys likely acted as corridors for postglacial expansion, longitudinal valleys, such as the Piave or Etsch valleys (indicated by ‘E’ and ‘P’, Figures 1C, 2B) did not facilitate postglacial expansions into the interior of the SLA. Given the species’ dispersal limitation, far-reaching postglacial expansion from refugial areas likely required an interconnected matrix of open habitats—specifically, forest-free valley floors or interconnected valley slopes (Figure 4). Notably, paleoecological evidence suggests that the temporal window in which such an open habitat matrix existed was short in the southern part of the SLA (R1, R2, Figure 1C), where forests advanced rapidly after the end of the LGM (Ravazzi et al. 2014). In contrast, the southeastern periphery of the SLA, including the outlined refugia of *A. italoaustriacus* in the periphery of the Tagliamento-Drau glacier system (R3, Figure 1C), remained forest-free much longer and well into the early Holocene (∼11 ka) due to the cold and dry continental climate in this area (Fritz 1967, Fritz 1972, Caf et al. 2023). We hypothesize that this delayed forest expansion may be key in enabling *A*. *italoaustriacus* postglacial colonization of interior parts of the Alps along transversal valleys from its refugium (Figure 4).

### Long-term survival of *A. italoaustriacus* despite glaciation-driven extinctions: The roles of microrefugial persistence and polyandry’s genetic benefits

The species *A*. *italoaustriacus* has its origins in the early Pleistocene (Calabrian) 1.78 ma, when the species diverged from its phylogenetic sister comprising *A*. *alpinus* and *A*. *ghilianii* (Figure 3A; relation to *A*. *ghilianii* not shown; Kirschner et al., in prep.). In contrast to its origin in the early Pleistocene, intraspecific diversification occurred significantly later (∼230 ka; Figure 3A). While we elaborate in detail how recurrent isolation linked to glaciation cycles may have steered lineage formation within *A*. *italoaustriacus* (Figure 3C, Figure 4), it is stunning that lineages older than 230 ka do not exist (Figure 3A). This is particularly interesting, as glaciation cycles from the mid-Pleistocene transition onwards (1,250 to 750 ka) were comparable in their intensity and duration (Herbert, 2023). This raises the question on why previous glaciations have not similarly produced lineages that survived until today. We speculate that the evolutionary past of *A*. *italoaustriacus* involved recurring and large-scale lineage extinctions, probably since the species’ very origin.

Despite undergoing repeated fragmentations into small populations and experiencing recurrent population bottlenecks, *A. italoaustriacus* has demonstrated remarkable resilience in avoiding complete extinction. In theory, the aforementioned processes diminish genetic diversity, lead to inbreeding, decrease population fitness and eventually lead to a population’s collapse (Charlesworth & Charlesworth, 1987; Gilpin & Soule, 1986). In case of *A*. *italoaustriacus*, glacial advances did not only repeatedly disrupt the species’ habitat, but have led to significant population decline, and may have also annihilated populations by trapping them on peaks and mountain chains, in particular in interior parts of the Alps (Figure 4). This is well reflected by the elaborated young age of populations in the ‘Interior Group’ (‘Schlern’ and ‘Peitler’, Figure 3A), and the modelled decrease in effective population size coinciding the LGM (Figure 3B). We conclude that this means that the long-term survival of the species has repeatedly depended on very few refugial populations ever since.

We propose that the species’ long-term resilience to extinction extends beyond mere “survivor’s luck” and reflects its ability to persist in microrefugia. Intrinsic traits favoring microrefugial survival have been discussed, and the ability to sustain high genetic diversity despite small population sizes has been highlighted as a key (Mosblech et al., 2011). Although polyandry—the reproductive strategy observed in all extant *Anonconotus* species (Vahed, 2002; Vahed & Carron, 2008)—has, to our knowledge, not been explicitly identified as such a trait, it is noteworthy that polyandrous mating systems have been shown to increase and maintain genetic diversity even in small and newly founded populations (Holman & Kokko, 2013; Lewis et al., 2020). As such this reproductive strategy may counteract the deleterious effects of inbreeding and reduced effective population sizes (Cornell & Tregenza, 2007), which are common in small or bottlenecked populations, and might be a key trait favouring survival in microrefugia.

We hypothesize that polyandry was crucial for the long-term survival of small *A. italoaustriacus* populations, and ensured high genetic diversity despite small population sizes and recurring bottlenecks (Figure 3B). This hypothesis is supported by the observed lack of inbreeding in all but one population (F_IS_ < 0, Table 1), as well as the higher-than-expected genetic diversity, evidenced by excess observed heterozygosity in all studied populations (ObsHet > ExpHet; Table 1). While this evidence may not be conclusive, we want to highlight that polyandry might be an overlooked but beneficial trait for survival in microrefugia specifically, and for long-term survival in high-altitude habitats in general. To our knowledge we are one of the first studies establishing this link. A systematic investigation into the prevalence of polyandrous mating in arthropods confined to high-altitude habitats and sky islands could provide valuable insights. Such research could broaden our understanding of how polyandry may influence genetic diversity in small, isolated populations confined to island-like habitats.

### Conservation of *A. italoaustriacus* should focus on stable refugia, monitoring population dynamics, and mitigation of human impact

The evolutionary history of *A. italoaustriacus* reveals both long-term survival in distributional stasis. Comparatively small areas, such as the Vette Feltrine, have served as long-term stable refugia, buffering populations from extreme climatic fluctuations across multiple glacial cycles. In addition to this, unique genetic variation has also evolved within isolated populations at the eastern range margin (‘Poella’, Figure 2). These findings highlight the critical importance of even very small areas, such as a single mountain range, for the long-term survival and conservation of high-altitude species with complex refugial histories (Mosblech et al., 2011; Tribsch, 2004).

We conclude that to conserve the genetic diversity of high-altitude species, efforts should prioritize both long-term stable refugia, such as the Vette Feltrine, and areas that harbor populations with younger and more dynamic evolutionary histories. Fortunately, many parts of the range of *A. italoaustriacus* are located within protected areas (e.g. population ‘Vette’: Parco Nazionale Dolomiti Bellunesi; populations ‘Schlern’, ‘Peitler’, ‘Poella’: Natura 2000 sites). However, apart from the Vette Feltrine, where the population biology of *A. italoaustriacus* is being studied in detail, also with the aim to inform local conservation management (Buzzetti, pers. comm., 2024), no similar efforts have been undertaken for the remaining populations. Expanding such research could be highly valuable, as data on census size, sex ratios, and other population metrics could be combined with genetic data to monitor the long-term effects of small population sizes and isolation.

Additionally, populations affected by human activities, such as touristic development (e.g., ‘Schlern’), would greatly benefit from close monitoring. This is particularly urgent given the species’ vulnerability to human impact evidenced by the likely human-induced eradication of a somewhat enigmatic population of *A. italoaustriacus* in the last century (Marangoni et al., 2020).

## Acknowledgements

We thank Filippo Maria Buzzetti from the Museo Civico di Rovereto (Rovereto, Italy) for sharing occurrence data and his valuable knowledge about the species. We are also grateful to Karim Vahed for inspiring discussions on the reproductive behaviour of *Anonconotus* bush-crickets. We acknowledge the conservation authorities of the provinces Bolzano/Bozen (Italy; general sampling permit issued for Petra Kranebitter and the Museum of Nature South Tyrol), Carinthia (Austria; Permit ID: SP3-NS-4298/2024), and the administration of the Parco Nazionale Alpi Bellunesi (Italy, Protocollo N. 4387/2020 del 02-09-2020) for issuing sampling permits. We thank Dominik Kirschner and Federico Marangoni for their assistance and companionship during field sampling. The computational results presented here have been achieved (in part) using the LEO high performance computing infrastructure of the University of Innsbruck (Innsbruck, Austria). This study was conducted in course of the project “Next Generation – Alpine Heuschrecken im Spannungsfeld von Isolation und Klimawandel” (CUP: H34I19000380005) funded by the “Forschungsfond der Südtiroler Landesmuseen” (Bolzano/Bozen, Italy).

## Data availability statement

Genomic sequence data generated through genotyping-by-sequencing is available in the NCBI Short Read Archive under BioProject PRJNA1292412. The mitochondrial sequence data (16S, cytochrome b) used to calibrate phylogenetic analyses will be published in a separate study (Kirschner et al., in prep.) focusing on the evolution of the genus *Anonconotus* and are available upon request.

