## Supplementary Material for "Glacial Legacies – How refugial dynamics shaped the evolution of the Alpine endemic bush-cricket *Anonconotus italoaustriacus*"

**Divergence time estimation based on mitochondrial genes**

In addition to the presented SNP-based coalescent tree (Figure 3) that was age calibrated using estimates from a phylogeny based on two mitochondrial genes (16s, Cytochrome b) and including all extant *Anonconotus* species (Kirschner et al. unpubl.), an independent rate-calibrated dating analysis was done to test robustness of node age estimates. This independent analysis was done on the basis of the partial sequences of four mitochondrial genes that are publicly available from Genbank (Martinez-Sañudo et al., 2022; Genbank accession numbers: OL364192.2 to OL364215.2). Specifically, these sequences represent parts of the mitochondrial Cytochrome Oxidase I (COI) and NADH dehydrogenase genes, and parts of the mitochondrial control regions 1 and 2 (CR1 and CR2). The dataset comprised a total of 24 individuals of *A*. *italoaustriacus* from four populations, of which three populations were sampled independently from the same localities as populations in this study. *A*. *alpinus* was used as an outgroup (accession numbers: OL470896.1, OL471011.1, OL580778.1, OL580782.1). All analyses were done in Beast v. 2.7.8 (Bouckaert et al., 2014). Input files were prepared in Beauti v. 2.7.8 (Bouckaert et al., 2014), using linked tree, and unlinked site and clock models. Unlike in Martinez-Sañudo et al. (2022), independent clock rates were used for each of the four genes accounting for rate heterogeneity. The HKY model was identified as the best substitution model for all genes using smart model selection (Lefort et al., 2017). An additional test for rate correlation (Tao et al., 2019) indicated autocorrelated rates in COI and NAD4 genes, and independent rates in CR1 and CR2. As a consequence, optimized relaxed clock models were used for COI and NAD4, and random local clock models for CR1 and CR2. This was not done by Martinez-Sañudo et al. (2022), who concatenated the four genes and treated them as a single sequence using a random local clock model. A uniform clock prior of 0.0125 (lower boundary: 0.1, upper boundary: 0.015) was assigned to the COI gene, corresponding to the mean clock rate of 0.016 substitutions per site per Ma suggested for COI in arthropods (Papadopoulou et al., 2010). Clock rates of the three remaining genes were set to be estimated by the Markov chain Monte Carlo (MCMC), and four independent MCMC chains were run for 50 million generations each. Resulting trees were combined using logcombiner v. 2.7.3 (Bouckaert et al., 2014) and treeannotator v. 2.7.3 (Bouckaert et al., 2014), whereas the first 25% of the trees were discarded as burnin. Tracer v. 1.6 (Rambaut et al., 2018) was used to inspect each run based on the criteria described for the coalescent analyses in Materials and Methods section. Trees from all four runs were combined using logcombiner and summarised into a maximum clade credibility tree using treeannotator. The resulting phylogenetic tree is shown in Supplementary Figure 6.

**Supplementary Table 1** Model evaluation and parameter estimates for all 2-population and 3-population models included in demographic model selection. AIC, Akaike information criterion; ωi, Akaike weight; theta, effective mutation rate of the reference population; nuA, ancestral population size; nu*, effective population size parameters; m*, migration rate parameters, T*, time parameters; s, fraction of ancestral population that founded derived population; f, fraction of updated population 2 to be derived from population 1. Graphical representations of all models are shown in Figure S1.

| **2-population model** | | |  |  |  |  |  |  |  |  |  |  |  |  |  |  |  |  |  |  |
| --- | --- | --- | --- | --- | --- | --- | --- | --- | --- | --- | --- | --- | --- | --- | --- | --- | --- | --- | --- | --- |
| **Allele numbers** | **Sum of SFS** | **Model** | **log-**  **likelihood** | **AIC** | **ωi** | **chi²** | **theta** | **nu2** | **m** | **m12** | **m21** | **T** | **T1** | **T2** | **s** | **f** |  |  |  |  |
| 8 / 22 | 2711.78 | founder nomig | -974.11 | 1954.22 | **0.61** | 4742.61 | 527.46 |  |  |  | 14.14 |  | 0.21 |  | 0.10 |  |  |  |  |  |
|  |  | founder nomig admix early | -974.15 | 1956.3 | 0.22 | 4730.89 | 527.95 | 15.08 |  |  |  | 0.21 |  |  | 0.10 | 0.01 |  |  |  |  |
|  |  | founder sym | -974.49 | 1956.98 | 0.15 | 4769.54 | 528.14 | 13.73 | 0.01 |  |  | 0.21 |  |  | 0.10 |  |  |  |  |  |
|  |  | founder asym | -975.72 | 1961.44 | 0.02 | 4797.68 | 523.52 | 13.86 |  | 0.02 | 0.04 | 0.22 |  |  | 0.10 |  |  |  |  |  |
|  |  | founder nomig admix late | -978.68 | 1965.36 | 0.00 | 4830.02 | 527.96 | 12.62 |  |  |  | 0.21 |  |  | 0.10 | 0.01 |  |  |  |  |
|  |  | founder nomig admix two epoch | -1078.78 | 2167.56 | 0.00 | 4482.39 | 625.64 | 1.09 |  |  |  |  | 0.18 | 0.21 | 0.50 | 0.98 |  |  |  |  |
|  |  | vic two epoch admix | -1142.97 | 2293.94 | 0.00 | 4175.77 | 837.86 |  |  |  |  |  | 0.40 | 0.16 | 0.50 | 0.98 |  |  |  |  |
|  |  | vic anc asym mig | -1186.46 | 2382.92 | 0.00 | 4434.53 | 805.47 |  |  | 9.89 | 0.22 |  | 0.57 | 0.10 | 0.50 |  |  |  |  |  |
|  |  | vic no mig admix late | -1208 | 2422 | 0.00 | 4397.9 | 675.01 |  |  |  |  | 0.17 |  |  | 0.50 | 0.01 |  |  |  |  |
|  |  | vic no mig | -1209.72 | 2423.44 | 0.00 | 4315.41 | 676.89 |  |  |  |  | 0.16 |  |  | 0.50 |  |  |  |  |  |
|  |  | vic no mig admix early | -1209.71 | 2425.42 | 0.00 | 4328.24 | 676.83 |  |  |  |  | 0.17 |  |  | 0.50 | 0.23 |  |  |  |  |
|  |  | vic anc sym mig | -1209.36 | 2426.72 | 0.00 | 4322.35 | 674.48 |  | 7.06 |  |  |  | 0.07 | 0.13 | 0.50 |  |  |  |  |  |
|  |  | vic sec contact asym mig | -1213.86 | 2437.72 | 0.00 | 4350.46 | 676.78 |  |  | 0.36 | 0.15 |  | 0.07 | 0.10 | 0.50 |  |  |  |  |  |
|  |  | vic sec contact sym mig | -1265.59 | 2539.18 | 0.00 | 4726.11 | 671.87 |  | 1.04 |  |  |  | 0.04 | 0.22 | 0.49 |  |  |  |  |  |
| **3-population model** | | |  |  |  |  |  |  |  |  |  |  |  |  |  |  |  |  |  |  |
| **Allele numbers** | **Sum of SFS** | **Model** | **log-**  **likelihood** | **AIC** | **ωi** | **chi²** | **theta** | **nu1** | **nuA** | **nu2** | **nu3** | **mA** | **m32** | **m31** | **m2** | **m3** | **T1** | **T2** | **T1b** | **T1a** |
| 8 / 8 / 8 | 4625.79 | split uni mig adjacent var3 | -1624.93 | 3265.86 | 1.00 | 14801.04 | 1102.94 | 0.61 | 0.62 | 0.34 | 0.26 | 0.07 | 0.03 |  |  |  | 0.23 | 0.15 |  |  |
|  |  | split uni mig adjacent var2 | -1658.39 | 3332.78 | 0.00 | 15544.44 | 897.84 | 0.67 | 0.43 | 0.49 | 0.43 | 1.11 |  | 0.08 |  |  | 6.58 | 0.23 |  |  |
|  |  | split uni mig adjacent var1 | -1670.94 | 3359.88 | 0.00 | 16580 | 694.42 | 0.85 | 0.75 | 0.58 | 0.41 | 0.65 | 0.50 | 0.03 |  |  | 3.02 | 0.26 |  |  |
|  |  | split sym mig adjacent var1 | -1691.17 | 3400.34 | 0.00 | 14286.71 | 1146.68 | 0.50 | 0.10 | 0.54 | 0.46 | 1.71 |  |  | 0.11 | 0.09 | 0.05 | 0.25 |  |  |
|  |  | refugia adj 2 var uni | -1721.74 | 3459.48 | 0.00 | 14477.15 | 1219.29 | 0.55 | 0.91 | 0.27 | 0.10 |  | 5.30 | 0.36 |  |  | 0.15 | 0.18 |  |  |
|  |  | split sym mig adjacent var2 | -1743.4 | 3502.8 | 0.00 | 23227.29 | 822.31 | 1.00 | 0.70 | 0.76 | 0.53 | 0.08 |  |  |  | 0.07 | 0.41 | 0.30 |  |  |
|  |  | refugia adj 3 var sym | -1843.72 | 3707.44 | 0.00 | 13594.14 | 1095.44 | 0.50 | 0.55 | 0.18 | 0.13 | 0.98 |  |  | 0.37 | 0.39 |  | 0.09 | 0.80 | 0.52 |
|  |  | refugia adj 3 var uni | -1936.76 | 3893.52 | 0.00 | 25497.82 | 753.77 | 1.08 | 25.0 | 0.21 | 0.09 | 0.10 | 6.03 | 0.28 |  |  |  | 0.14 | 0.61 | 0.24 |
|  |  | refugia adj 2 var sym | -2177.62 | 4371.24 | 0.00 | 42511.51 | 674.76 | 1.14 | 1.70 | 0.83 | 0.22 |  |  |  | 0.47 | 0.47 | 0.95 | 0.44 |  |  |
|  |  | split sym mig adjacent var3 | -2606.52 | 5229.04 | 0.00 | 24191.69 | 938.71 | 0.89 | 0.31 | 1.78 | 0.80 | 0.16 |  |  |  | 0.62 | 0.24 | 0.21 |  |  |

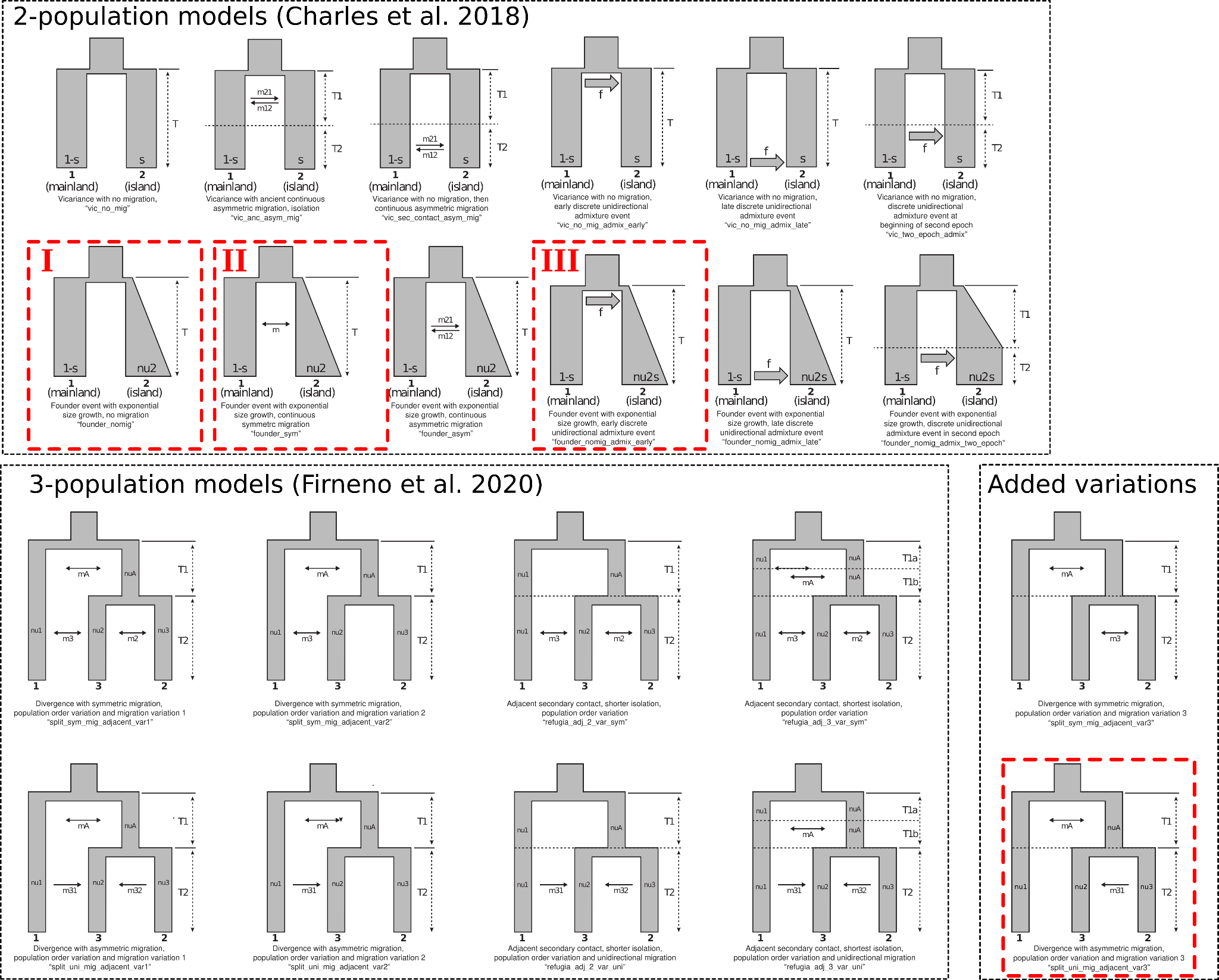

**Figure S1** Graphic representation of all 2-population and 3-population models used for demographic modeling (model definition as in Charles et al., 2018; Firneno et al., 2020). Best-performing models are outlined in red, with the best-performing 2-population models ranked by descending conditional likelihood (I–III; Figure 2, Table S1)

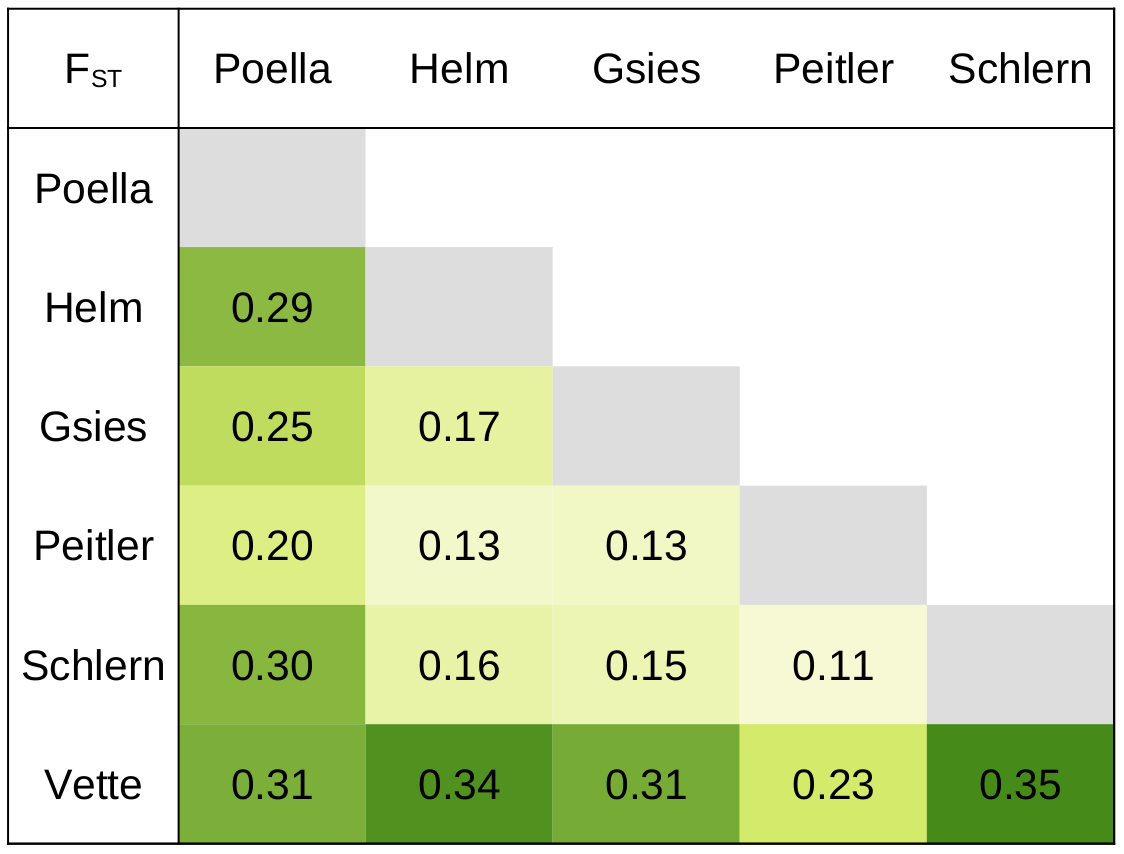

**Figure S2** Matrix of pairwise F_ST_ values among all populations.

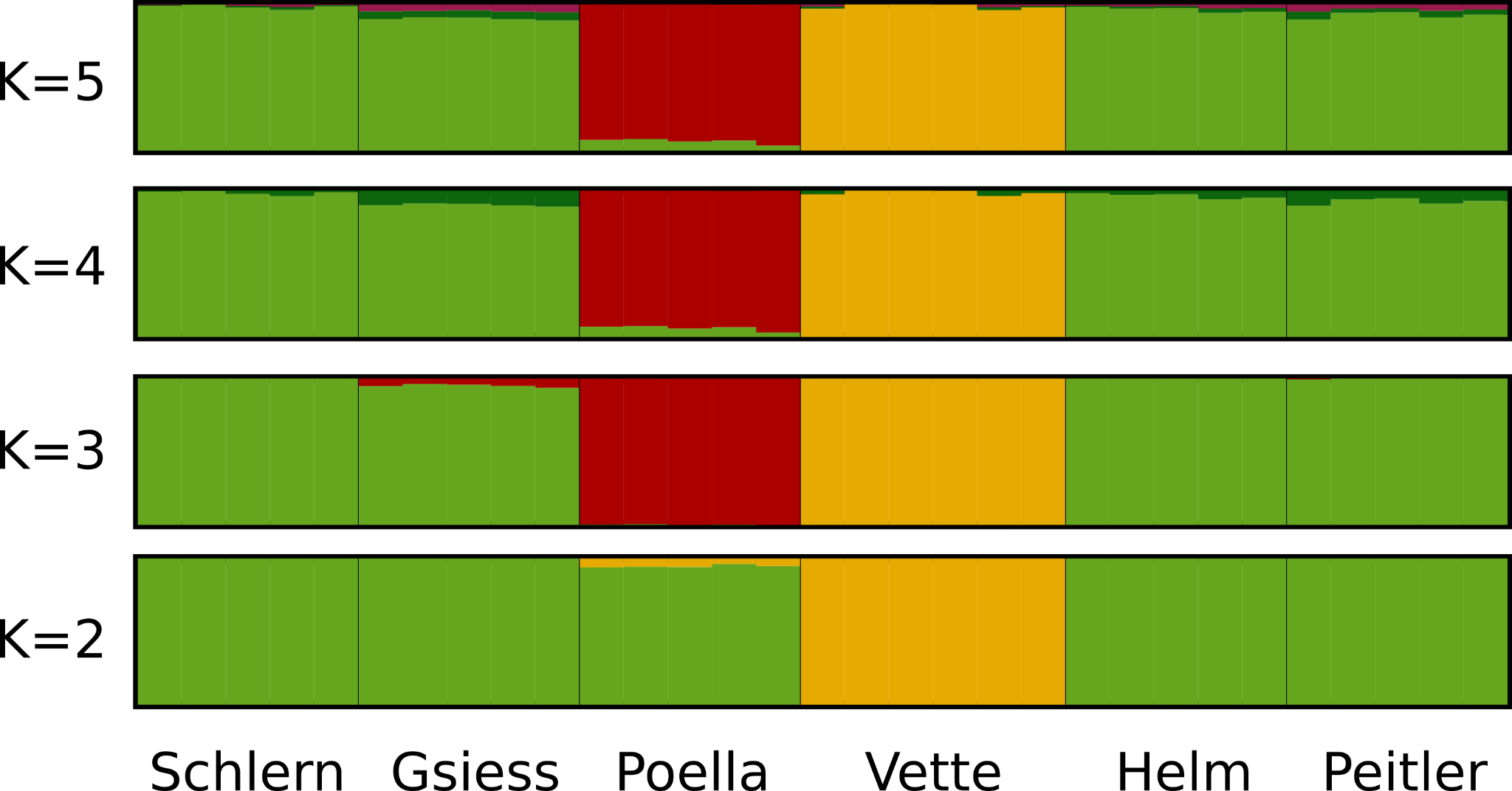

**Figure S3** Results from Bayesian genetic clustering assuming K=2–5 clusters. Each barplot represents one individual, populations are separated by black lines, and population names are indicated below. Colors indicate the proportion of variation assigned to each genetic cluster.
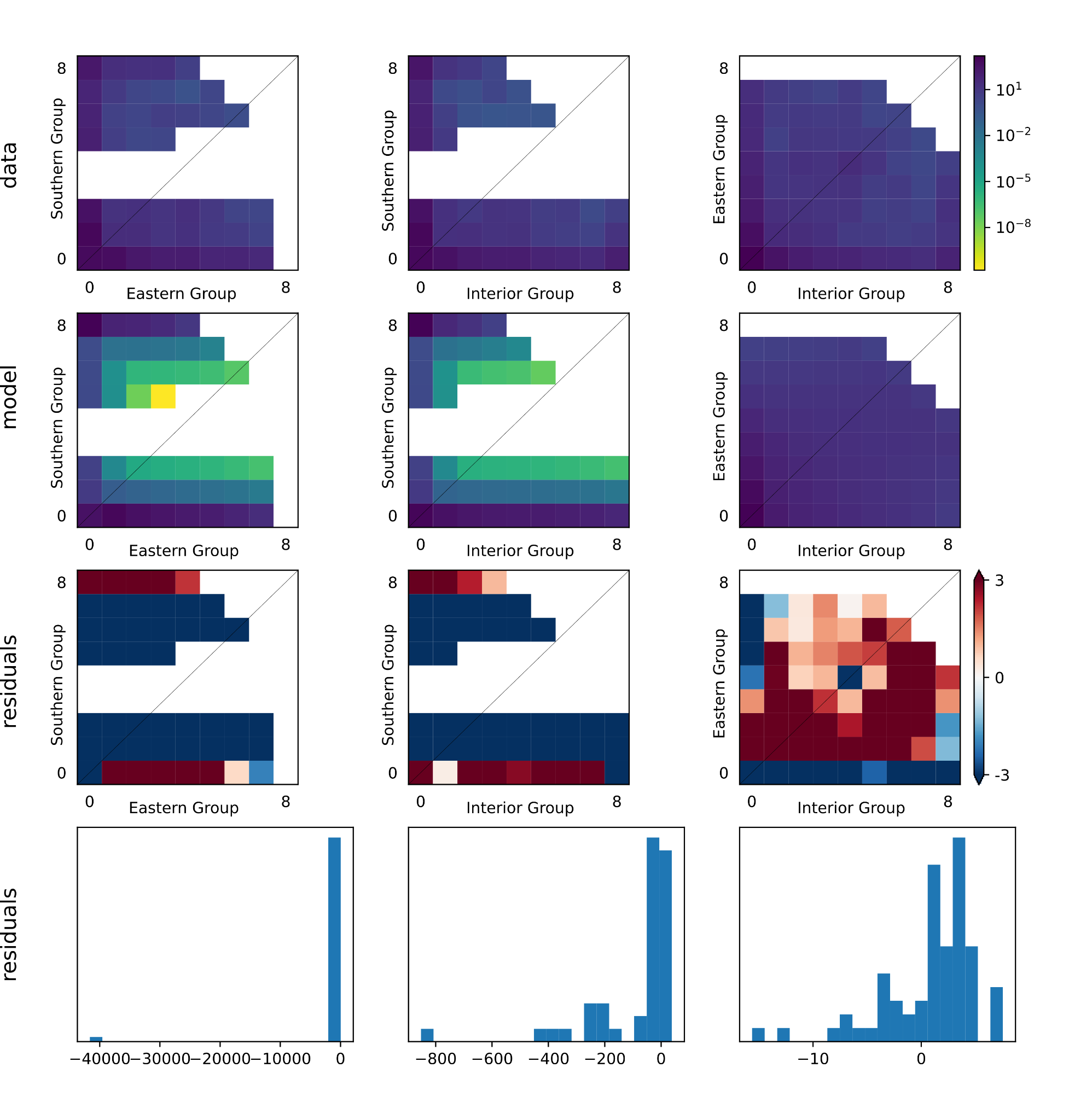

**Figure S4** Visual representation of the joint site frequency spectra inferred from empiric data and inferred for the best-fitting demographic model (3-population model ‘split_uni_mig_adjacent_var_3’; Figure S1), and the corresponding residuals.

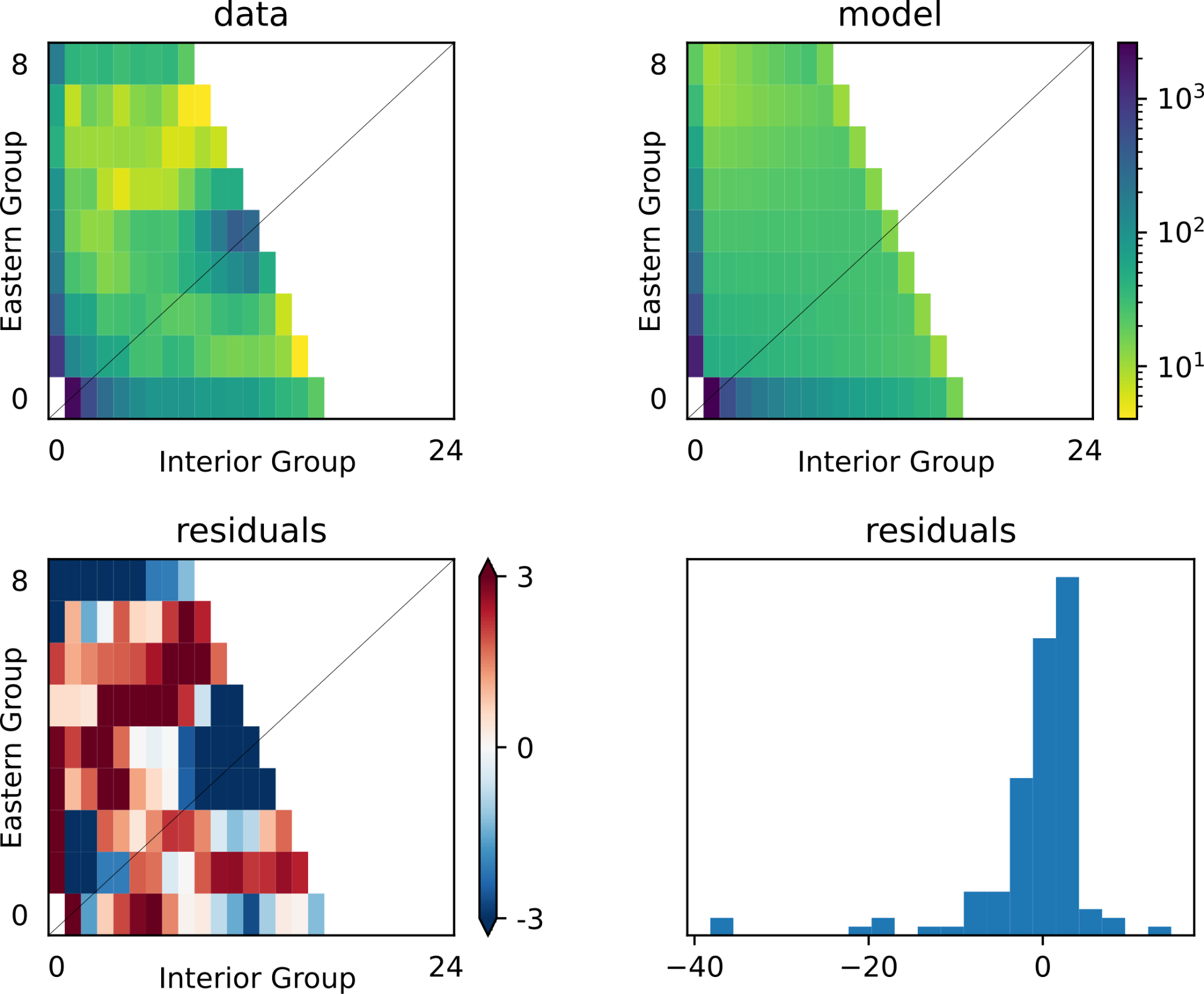

**Figure S5** Visual representation of the joint site frequency spectra inferred from empiric data and inferred for the best-fitting demographic model (2-population model ‘founder_no_mig’; Figure S1), and the corresponding residuals.

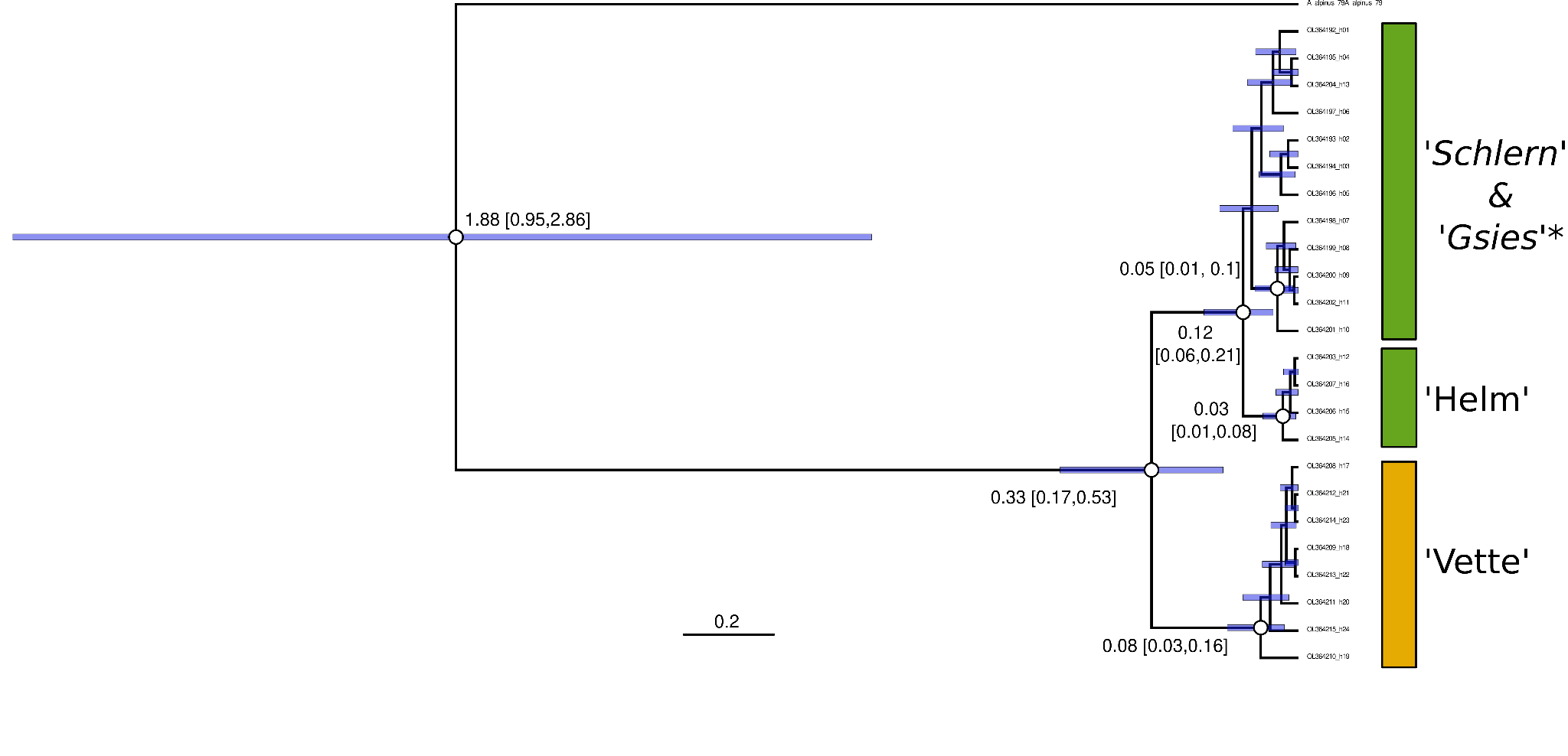

**Figure S6** Phylogenetic tree based on parts of four mitochondrial genes. Node labels represent rate-calibrated divergence time estimates, with 95% highest posterior density ranges shown in parentheses and by blue bars. White circles at nodes indicate posterior probabilities > 95%. Tip labels are according to the genbank accessions. Colored bars on the right indicate overlap of the used genbank accessions samples with the populations sampled and analyzed in this study.
